# How much do model organism phenotypes contribute to the computational identification of human disease genes?

**DOI:** 10.1101/2021.12.24.474099

**Authors:** Sarah M. Alghamdi, Paul N. Schofield, Robert Hoehndorf

**Affiliations:** Computational Bioscience Research Center (CBRC), King Abdullah University of Science and Technology, 4700 KAUST, 23955 Thuwal, Saudi Arabia; Department of Physiology, Development & Neuroscience, University of Cambridge, Downing Street, CB2 3EG, Cambridge,United Kingdom

**Keywords:** model organism, phenotype, disease gene discovery, ontology, semantic similarity, machine learning

## Abstract

Computing phenotypic similarity has been shown to be useful in identification of new disease genes and for rare disease diagnostic support. Genotype–phenotype data from orthologous genes in model organisms can compensate for lack of human data to greatly increase genome coverage. Work over the past decade has demonstrated the power of cross-species phenotype comparisons, and several cross-species phenotype ontologies have been developed for this purpose. The relative contribution of different model organisms to identifying disease-associated genes using computational approaches is not yet fully explored. We use methods based on phenotype ontologies to semantically relate phenotypes resulting from loss-of-function mutations in different model organisms to disease-associated phenotypes in humans. Semantic machine learning methods are used to measure how much different model organisms contribute to the identification of known human gene–disease associations. We find that only mouse phenotypes can accurately predict human gene–disease associations. Our work has implications for the future development of integrated phenotype ontologies, as well as for the use of model organism phenotypes in human genetic variant interpretation.

## INTRODUCTION

Discovering and building models of human phenotypes in nonhuman animals has, over the last half century, proved to be of substantial importance in improving our understanding of human disease and its underlying biology (Aitman et al., 2011; Wangler et al., 2017; Brown, 2021; Baldridge et al., 2021), and is providing insights that may be used to develop new therapeutic and diagnostic capabilities. The amount of data available which relates genetics, in particular genomic variation, to phenotypes associated with disease, is increasing rapidly. For example, the Monarch Initiative lists more than 2M phenotypic associations over more than 100 species from dozens of public resources (Shefchek et al., 2020). By comparing the similarities between phenotypic profiles this data can be used to help understand gene function and to identify the genotypic origins of phenotypic variation, which has wide applications in the discovery of the etiology of disease and the identification of candidate disease genes.

The challenge of relating phenotypes accross different species is very significant. The ontologies and controlled vocabularies used to describe phenotypes are species-specific and often structured in markedly different ways (Gkoutos et al., 2017). In order to compare phenotypic profiles between species, several different approaches have been developed to create an overarching phenotype ontology allowing the integration of phenotype-genotype data from multiple species. This can then be used for measuring phenotypic similarity between an instance of one species, for example a human with a genetic disorder, and phenotypes annotated to multiple species and genotypes. This approach mobilises the huge amount of genotype-phenotype data available in public databases such as Mouse Genome Informatics (MGI) (Eppig et al., 2017; Ringwald et al., 2021), Flybase (Larkin et al., 2020) and Online Mendelian Inheritance in Man (OMIM) (Amberger and Hamosh, 2017), and maximizes the possibility of finding a phenotype annotation to a potential disease gene where such a phenotype has not yet been reported in humans.

The development of a phenotype ontology covering both humans and model organisms has been essential to this task. The main approaches use evolutionary homology (and analogy) between anatomical structures (Mungall et al., 2012) and physiological processes, formalize these in a knowledge base or ontology, and infer relations between phenotypes using automated reasoning (Matentzoglu, Osumi-Sutherland, Balhoff, Bello, Bradford, Cardmody, Grove, Harris, Nomi Harris, Köhler, McMurry, Mungall, Munoz-Torres, Pilgrim, Robb, Robinson, Segerdell, Vasilevsky and Haendel, 2019; Hoehndorf et al., 2011).

Loss-of-function phenotypes are available in several model organisms through hypothesis-driven and large-scale reverse genetics experiments (Brown et al., 2018; Peterson and Murray, 2021), and genotype–phenotype data from model organisms, combined with cross-species phenotype ontologies, has been used to discover human disease-associated genes using measures of phenotype similarity (Smedley and Robinson, 2015; Meehan et al., 2017; Hoehndorf et al., 2011). The underlying assumption of phenotype-based methods to discover disease-associated genes is that genes function in evolutionary conserved pathways or modules, and phenotypes associated with a loss or change of function in a gene, are similar to phenotypes observed in a loss or change of function in the human ortholog of that gene (Oti and Brunner, 2007; McGary et al., 2010; Oti et al., 2008; Barabási et al., 2010). These methods are not only used to identify disease-associated genes but also to interpret and prioritize genomic variants associated with disease in tools that combine variant pathogenicity prediction with ranking of candidate genes (Cipriani et al., 2020; Boudellioua et al., 2017).

Phenotype-based methods to identify candidate genes associated with a set of phenotypes in humans are highly successful when the human gene has already been identified as a disease gene and is therefore associated with phenotypes (Köhler et al., 2009), and there are many examples where mouse phenotypes closely resemble human phenotypes and have therefore been used to identify disease-associated genes in humans (Meehan et al., 2017; Brommage et al., 2019; Smedley et al., 2021). Identification of candidate Mendelian disease genes using high-throughput screening suggests that this strategy might be able to identify candidates for inherited diseases of unknown genetic etiology. For example, out of 3,328 genes screened in the mouse, potential models for 360 diseases were reported including novel candidates (Meehan et al., 2017). More recently, IMPC reported knockouts of 1,484 known disease genes, approximately half of which showed phenotypic similarity to human diseases using the Phenodigm platform (Cacheiro et al., 2019). It is estimated that of the 16,847 mouse genes with a human ortholog, 79.9% have a null allele, either derived from hypothesis-driven experiments or large-scale screens such as the IMPC (Peterson and Murray, 2021); there are currently 3,381 genes with mouse–human orthologs for which there are no corresponding mouse loss-of-function phenotypes. MGI reports 1,694 Human diseases with one or more mouse models and 7,142 mouse genotypes modeling human diseases (MGI version 6.17; 14 December 2021), but their interpretation is complicated by the inclusion of dominant inheritance and multigenic or humanized models. It has been suggested that the “phenotype gap” might be filled with genotype-phenotype associations from non-mammalian organisms with complementary coverage to the mouse and where loss-of-function mutations in mouse–human orthologs have no phenotype data (Mungall et al., 2016). To date, the contribution of different model organisms to the computational phenotype-driven identification of human disease genes has not been critically evaluated, an assessment that is important for the continued development of strategies and computational approaches to disease gene discovery. It is important to understand and quantify the contribution of more distant model organisms to discovering human disease-associated genes using the methods that have so successfully been applied to the mouse, in particular as, for example, zebrafish phenotypes are used in methods for disease gene discovery and human genetic variant interpretation (Wangler et al., 2017; Smedley et al., 2015; 2016).

We use two different cross-species ontologies and several state of the art methods for phenotype-based identification of disease-associated genes to evaluate the contribution of mouse, zebrafish, fruitfly, and fission yeast loss-of-function phenotypes to discovering human disease genes. We find that only the mouse consistently predicts disease genes whereas the organisms that are more distant do not contribute. As part of our analysis, we find that our evaluation is affected by several biases in how orthologs of disease-associated genes are annotated in model organism databases as well as how phenotype-based methods exploit these annotations; we analyze and correct for some of these biases to support future work in relating phenotype data to human disease.

## RESULTS

### Contribution of model organisms to disease gene discovery

We collected phenotypes associated with loss-of-function mutations in the mouse, zebrafish, fruitfly, and fission yeast, from model organism databases. The phenotypes are described using different organism-specific phenotype ontologies and we combine the phenotypes using the integrated phenotype ontologies uPheno (Shefchek et al., 2020) and our extension of the PhenomeNET ontology (Pheno-e). Both phenotype ontologies combine the classes that represent phenotypes in different model organisms within a single ontology, thereby allowing us to exploit relations between the phenotypes and compare them. Pheno-e and uPheno also include human phenotypes from the Human Phenotype Ontology (HPO) (Köhler et al., 2021) thereby allowing us to relate mutant model organism phenotypes to human disease-associated phenotypes.

We used the Pheno-e and uPheno ontologies and the phenotypes associated with loss-of-function mutations and human Mendelian diseases to test whether, and how much, different model organisms contribute to the phenotype-based computational discovery of disease-associated genes. For the purpose of evaluating the predictive performance, we used two datasets of gene–disease association: a “human” dataset which includes associations of human genes with Mendelian diseases reported in the Online Mendelian Inheritance in Man (OMIM) (*Online Mendelian Inheritance in Man (OMIM)*, 2020) database, and a “mouse” evaluation set which consists of associations of mouse genes with human disease and represents mouse models of human disease in the MGI database (Ringwald et al., 2021). Then, we measure the semantic similarity between the phenotypes resulting from a gene’s loss of function and human diseases (see Figure S1). For each disease, we rank all genes by their phenotypic similarity to the disease; we then determine at which rank we identify orthologs of known disease-associated genes.

This approach has repeatedly been successfully applied to discover disease-associated genes from model organisms through ontology–based computation of phenotype similarity (Meehan et al., 2017; Washington et al., 2009; Smedley et al., 2021), and further forms the foundation of several computational methods for finding disease-associated genomic variants (Smedley and Robinson, 2015; Smedley et al., 2016; Boudellioua et al., 2017). Multiple different approaches for determining phenotypic similarity have been developed, ranging from hand-crafted semantic similarity measures (Köhler et al., 2009; Smedley et al., 2013; Pesquita et al., 2009) to machine learning approaches (Smaili et al., 2018*a*; Chen et al., 2020). We used four different approaches to compute phenotype similarity between model organism phenotypes and human disease. First, we use Resnik’s semantic similarity measure (Resnik, 1999) which relies on the taxonomic relations in the phenotype ontology to determine similarity between two sets of phenotypes. Resnik’s similarity compares two phenotype classes whereas we need to compare two sets of phenotype classes (i.e., all the phenotypes associated with the disease and all the phenotypes observed in the model organism). Consequently, we use the “best match average” strategy (Pesquita et al., 2009) (see Materials & Methods) to combine multiple pairwise similarity measurements into a similarity between two sets of phenotypes. Resnik’s similarity uses only the ontology taxonomy whereas phenotype ontologies contain a large amount of additional information in the form of axioms that provide a computational description of the intended meaning of phenotype classes (Hoehndorf et al., 2015; Gkoutos et al., 2017). Therefore, we use the unsupervised machine learning method OPA2Vec (Smaili et al., 2018*a*) which is a deep learning method that learns a “representation” of sets of phenotypes based on ontology axioms as well as natural language information contained in ontologies such as labels and definitions. As a third and fourth approach, we use the deep learning methods OWL2Vec* (Chen et al., 2021) and DL2Vec (Chen et al., 2020) which first converts ontology axioms into a graph, applies a random walk to explore the neighborhood of nodes in that graph, and then generates a feature vector using Word2Vec. The aim of using these methods based on random walks is to exploit more “distant” relations that arise through connecting multiple ontology classes. Figure 1 illustrates the different approaches.

**Fig. 1.**
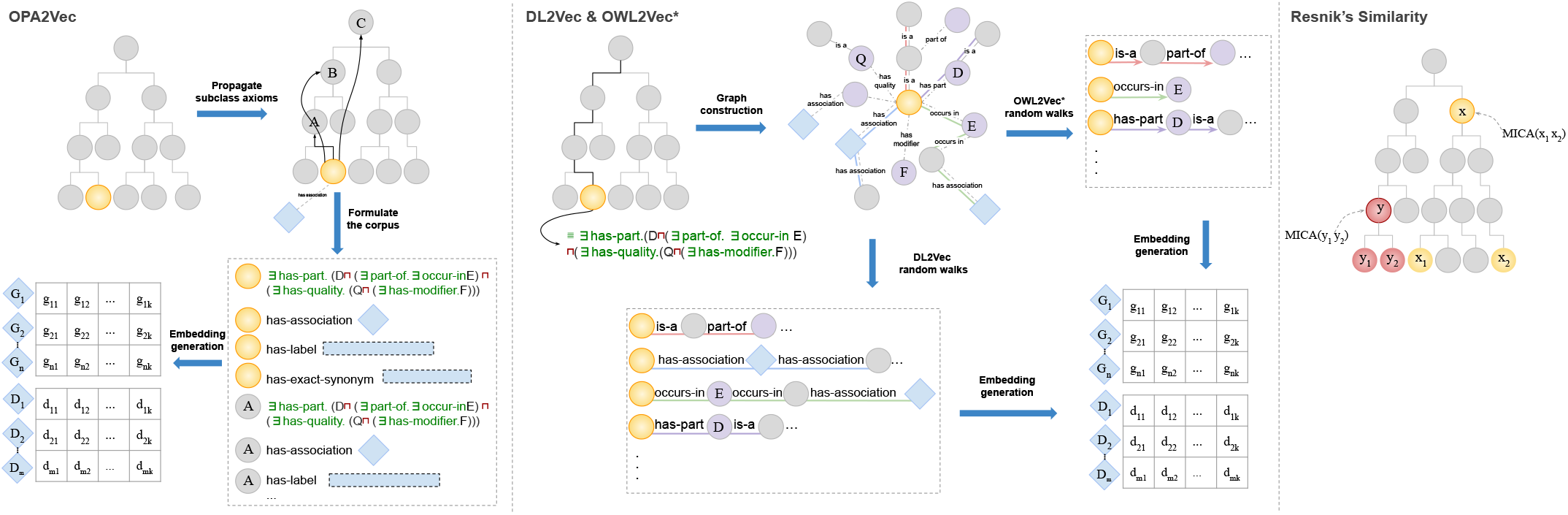
Illustration of the approaches that we used to calculate phenotypic similarity. Resnik’s similarity uses the taxonomy of the ontology. DL2Vec and OWL2Vec generate a graph from the ontologies axioms then perform random walks to generate vector representations for genes and diseases, with some differences including that the graphs are directed in OWL2Vec and undirected in DL2Vec. OPA2Vec generates vector representations by using the axioms of the ontologies propagated over the subsumption hierarchy along with the natural language information available in the ontology.

In our first experiment, we focus only on the groups of orthologous genes that have phenotype annotations in the mouse, zebrafish, fruitfly, and fission yeast; the aim is to compare the contributions of different model organism to discovering gene–disease associations on the same set of associations from the “human” dataset. There are 255 human genes with orthologous genes annotated with phenotypes in all organisms we consider, and of these, 88 have a human ortholog associated with a Mendelian disease; several genes are associated with more than one Mendelian disease, and, in total, the 88 genes are associated with 173 Mendelian diseases.

We compare the phenotypic similarity of these genes to human disease phenotypes and, within each organism, we rank the genes by their similarity to each disease. We then evaluate the ranks at which we discover the “correct” gene (i.e., the gene with the human ortholog that is associated with the disease) and quantify the results using the ROCAUC measure (see Materials & Methods). Table 1 summarizes the resulting performance. The results indicate that mouse mutant phenotypes can be used to reliably detect human disease-associated genes by all methods, whereas the other organisms do not consistently show a positive signal, and the quality of the signal is very dependent on the method used.

**Table 1.** Comparison of the performance of predicting gene-disease associations evaluated on diseases associated with genes which have orthologs with at least one phenotype annotation in mouse, fish, fly, and yeast (255 genes). Results are bold if they are significantly different from random (i.e., the confidence intervals does not overlap with the ROCAUC 0.5 of a random classifier).

| Overlapping Dataset | Mouse | Fish | Fly | Yeast |
| --- | --- | --- | --- | --- |
| Resnik similarity | **0.663** ± 0.069 | 0.558 ± 0.072 | 0.572 ± 0.085 | 0.475 ± 0.069 |
| OPA2Vec | **0.599** ± 0.072 | **0.585** ± 0.072 | 0.526 ± 0.074 | 0.544 ± 0.069 |
| OWL2Vec* | **0.624** ± 0.071 | 0.479 ± 0.073 | 0.472 ± 0.076 | 0.483 ± 0.069 |
| DL2Vec | **0.607** ± 0.071 | 0.517 ± 0.073 | 0.486 ± 0.075 | **0.594** ± 0.068 |

However, our observations are based on a relatively small set of 88 disease-associated genes that have orthologs with phenotypes in all organisms we study. Therefore, we analyze all genes with phenotypes in the different model organisms separately, incorporating genes that may lack phenotype annotations in other model organisms. As in our first experiment, we determine the phenotypic similarity using different semantic similarity measures and evaluate how well-established associations can be recovered.

Table 2 summarizes the ROCAUC values for each organism using the four approaches (Resnik similarity, OPA2Vec, OWL2Vec*, and DL2Vec). Similar to the first experiment, mouse phenotypes show the highest performance across all methods we consider and the mouse is the only organism where all four methods to compute phenotype similarity show a predictive performance that is better than random. Resnik similarity shows better-than-random performance for zebrafish and fruitfly phenotypes, but other methods predict disease-associated genes no better than a random classifier (except DL2Vec in fission yeast using human gene–disease associations); in evaluations based on ontology embedding methods the predicted performance is even significantly “worse than random” (i.e., significantly below the ROCAUC 0.5 of a random classifier); this indicates that increased phenotypic dissimilarity between a gene and disease is associated with a higher chance of the gene and disease being associated, a rather counter-intuitive result that requires further exploration. We tested the hypothesis that these results are due to a study bias which results in an increased (phenotypic) distance due to the ontology structure. We break this hypothesis into two parts; first, we hypothesize that genes that have an ortholog that is associated with a Mendelian disease in humans have more, and more specific, phenotype annotations than genes whose ortholog is not associated with a Mendelian disease (or for which no human ortholog is known); this hypothesis tests for a form of study bias within the phenotype annotations. We find that disease-associated genes have a significantly higher total information content compared to non-disease associated genes (mouse: *p* = 1.361 · 10^*−*43^, fish: *p* = 6.793 · 10^*−*20^, fly: *p* = 1.115 · 10^*−*12^, yeast: *p* = 0.003; one-tailed *t*-test).

**Table 2.** Predicting gene-disease associations

| Human evaluations data set |  |  |  |  |
| --- | --- | --- | --- | --- |
|  | Mouse | Fish | Fly | Yeast |
| Resnik's | **0.786** ± 0.011 | **0.598** ± 0.018 | **0.595** ± 0.018 | 0.525 ± 0.032 |
| OPA2Vec | **0.630** ± 0.013 | 0.459 ± 0.017 | 0.306 ± 0.016 | 0.517 ± 0.032 |
| OWL2Vec* | **0.711** ± 0.012 | 0.449 ± 0.018 | 0.418 ± 0.018 | 0.500 ± 0.032 |
| DL2Vec | **0.761** ± 0.011 | 0.473 ± 0.018 | 0.389 ± 0.017 | **0.557** ± 0.032 |
| Mouse evaluations data set |  |  |  |  |
|  | Mouse | Fish | Fly | Yeast |
| Resnik's | **0.942** ± 0.009 | **0.592** ± 0.026 | **0.639** ± 0.039 | 0.525 ± 0.061 |
| OPA2Vec | **0.696** ± 0.017 | 0.462 ± 0.025 | 0.304 ± 0.035 | 0.406 ± 0.060 |
| OWL2Vec* | **0.817** ± 0.014 | 0.447 ± 0.026 | 0.435 ± 0.038 | 0.479 ± 0.061 |
| DL2Vec | **0.887** ± 0.012 | 0.460 ± 0.026 | 0.404 ± 0.037 | 0.447 ± 0.061 |

If these phenotypes do not match human disease-associated phenotypes well, the distance between these specific (i.e., “deep” within the ontology hierarchy) phenotypes and general (i.e., “shallow” within the ontology hierarchy) human phenotypes is higher than for less specific phenotypes; for example, the distance between the very general human phenotype class *Phenotypic abnormality* (`HP:0000118`) and the general fly phenotype *Phenotypic abnormality of organism* (`FBbtAB:00000001`) is less than the distance between *Phenotypic abnormality of organism* (`FBbtAB:00000001`) and the more specific class *Phenotypic abnormality of eye dorsal compartment* (`FBbtAB:00111608`). To further test whether this holds true across all genes with disease-associated and non-associated homologs in human, we calculate the absolute difference in information content between the phenotypes of the fly model and the most informative human phenotype superclass; the average difference in information content for genes with disease-associated human orthologs is 44 whereas the average difference in information content is 14 for genes with non-associated orthologs (*p* ≤ 1 · 10^*−*60^, Student’s t-test; see Supplementary Materials Section 2). The only method that is not based on distances in our test is Resnik’s similarity (which relies on the information content of the most informative shared ancestor), and this is also the only method not showing ROCAUCs below 0.5. Overall, these tests demonstrate that the ROCAUC results significantly lower than 0.5 are due to study bias combined with how the similarity methods utilize the ontology structure to determine similarity (i.e., based on distances traversed between classes).

As the mouse is the only model organism that consistently predicts gene–disease associations, we tested whether combining mouse phenotypes with other organism phenotypes would change the prediction results, i.e., whether combining information from multiple model organisms can improve predictions (i.e., test whether phenotypes of different organisms complement each other). We tested this on varying sets of genes depending on whether they have phenotypes in two model organisms. Table S1 shows the results. We find that combining mouse phenotypes with phenotypes of other model organisms does not significantly change prediction results.

So far, we performed our analysis only using the Pheno-e ontology. It was unclear whether our results demonstrated an inability of the Pheno-e ontology to compare phenotypes adequately or if they reflect a property of the underlying data and the methods used to analyze it. Consequently, we used the cross-species phenotype ontology uPheno (Shefchek et al., 2020) and repeated the same analysis of predicting gene–disease associations using the four phenotype similarity computation methods; the results and comparison to Pheno-e are shown in Table 3. The results indicate that Pheno-e and uPheno have comparable performance and do not consistently show significant differences in predictive performance across different model organism and analysis methods.

**Table 3.** Comparison of the performance of the Pheno-e and the uPheno ontologies to predict gene-disease associations using mouse, fly and yeast on the human evaluation data set. Note that Fish data are not included in this comparison as uPheno uses the zebrafish phenotype ontology and Pheno-e defines fish phenotype classes axiomatically

|  | Mouse |  | Fly |  | Yeast |  |
| --- | --- | --- | --- | --- | --- | --- |
|  | Pheno-e | uPheno | Pheno-e | uPheno | Pheno-e | uPheno |
| Resnik's | **0.786** ± 0.011 | **0.746** ± 0.011 | **0.593** ± 0.018 | **0.645** ± 0.018 | 0.525 ± 0.032 | 0.509 ± 0.032 |
| OPA2Vec | **0.626** ± 0.013 | **0.590** ± 0.013 | 0.375 ± 0.018 | 0.326 ± 0.017 | 0.522 ± 0.032 | 0.519 ± 0.032 |
| OWL2Vec | **0.711** ± 0.012 | **0.751** ± 0.011 | 0.418 ± 0.018 | 0.488 ± 0.018 | 0.500 ± 0.032 | 0.514 ± 0.032 |
| DL2Vec | **0.755** ± 0.011 | **0.756** ± 0.011 | 0.444 ± 0.018 | 0.436 ± 0.018 | **0.550** ± 0.032 | **0.538** ± 0.032 |

### Supervised prediction

One advantage of similarity measures that rely on embeddings is that they can be used as input to “supervised” machine learning approaches and thereby give rise to supervised similarity measures (Smaili et al., 2018*b*). In supervised machine learning, some examples of existing and absent associations between genes and diseases are used to train a model that can determine whether a new gene–disease pairs is associated or not. Using the ontology embeddings as input to supervised machine learning methods has previously resulted in significantly improved prediction of gene–disease associations (Smaili et al., 2018*a*).

We train a machine learning model (an artificial neural network), and use the output of this model to classify pairs of gene- and disease-embedding into two classes, depending on whether the gene is associated with the disease (positives) or not (negatives). We evaluate the performance using a 10-fold cross-validation strategy (see Materials & Methods); Table 4 shows the results. We find that the supervised machine learning approach improves the predictive performance significantly over the unsupervised similarity-based approach, not only when using mouse model phenotypes but also for all other organisms. Furthermore, the supervised model is able to predict gene–disease associations significantly better than a random classifier using all embedding methods and organisms, and further improves significantly over all unsupervised prediction approaches.

**Table 4.** Predicting gene-disease associations using supervised methods and our proposed naïve classifier

| Human evaluations data set |  |  |  |  |
| --- | --- | --- | --- | --- |
|  | Mouse | Fish | Fly | Yeast |
| MLP - OPA2Vec | 0.898 ± 0.007 | 0.836 ± 0.013 | 0.872 ± 0.012 | 0.763 ± 0.0275 |
| MLP - OWL2Vec | 0.875 ± 0.009 | 0.826 ± 0.013 | 0.874 ± 0.012 | 0.775 ± 0.027 |
| MLP - DL2Vec | 0.897 ± 0.007 | 0.880 ± 0.012 | 0.893 ± 0.011 | 0.781 ± 0.0267 |
| Naïve Classifier | 0.722 ± 0.011 | 0.689 ± 0.016 | 0.753 ± 0.014 | 0.510 ± 0.031 |

However, while the predictive performance is substantially higher than random, it is somewhat surprising that the predictive performance when using phenotypes from distant organisms such as fly or fish matches the performance of using mouse phenotypes, and that even yeast phenotypes apparently are able to identify a large number of gene–disease associations quite accurately when there are so few orthologous genes (estimated to be around 2,000 (O’Brien, 2004)), many without known disease associations in OMIM, the evaluation dataset. Neural networks may be able to exploit non-biological signals in training datasets to achieve relatively high predictive performance without producing biologically meaningful prediction results. For example, genes that are well studied and have a higher number of annotations may be associated with more diseases, or more likely be associated with diseases; ranking genes higher solely based on the number of annotations they received could therefore improve prediction performance even without a specific biological signal. To test this hypothesis, we design a “naïve” classifier that predicts gene–disease associations solely based on the sum of the information content of phenotypes within a gene, i.e., it can be used to test whether genes that are annotated with more and more specific phenotypes are more likely associated with any disease. The naïve classifier ranks all genes based on the sum of the information content of their phenotype annotations, and predicts, for each disease *D*, the genes in descending order ranked by their information content; this prediction is independent of the disease *D*, i.e., the same list of genes is predicted in the same order for each disease (see Materials & Methods). The result of the prediction by the machine learning model, together with the naïve classifier results, are shown in Table 4. The results demonstrate that there is substantial bias in the underlying data that can be exploited by the naïve classifier, and is likely exploited by the machine learning models as well.

## DISCUSSION

We have evaluated the contribution of different model organism phenotypes to the computational identification of human gene– disease associations through the use of a variety of semantic similarity and machine learning methods. We find that the main contribution towards discovering human disease-associated genes using these methods comes from mouse phenotypes, whereas other model organism data do not contribute significantly to this task. The premise that pooling genotype–phenotype data from multiple organisms to enhance the phenotype-driven prediction of human disease genes, or interpret human genetic variants, is in principle sound, and has driven the development of multiple cross-species phenotype ontologies. The assumption has been that, as long as the knowledge contained in the ontology is “true”, then this should help bridge the “phenotype gap”, i.e., the human genes that have no phenotype associations in human but do have in model organisms. However, a critical evaluation of the main types of methods in use, machine learning and semantic similarity, indicates that the contribution of the non-mammalian model organism phenotypes to this task is computationally insignificant in comparison to mouse data. We identify two problems with the inconsistency of the results obtained by different methods; the first is bias generated by the use of the structure of the cross-species ontologies available, and the second we have identified as issues such as annotation density; however, there may be further biases that affect the results.

We tested the impact of a number of different parameters on our finding. First, our results hold true across two cross-species phenotype ontologies, Pheno-e and uPheno (Matentzoglu, OsumiSutherland, Balhoff, Bello, Bradford, Cardmody, Grove, Harris, Harris, Köhler et al., 2019). Both ontologies have similar content and goals but are based on different ontology design patterns (Gkoutos et al., 2017; Alghamdi et al., 2019). We compared the two ontologies in our analyses to test whether the underlying ontology design patterns have a significant impact but we did not consistently identify significant differences between both ontologies, indicating that our results hold true independent of the choice of phenotype ontology. Further, we used different analysis methods, focusing both on traditional semantic similarity measures (Pesquita et al., 2009) that are largely defined based on explicit assumption of how similarity should be computed, as well as methods based on unsupervised and supervised machine learning with ontologies (Kulmanov et al., 2020). The machine learning methods we employed are largely based on “paths” in graph-based representations of the ontologies, whereas the semantic similarity we used is based on information content of classes without considering “paths” explicitly; in particular, “distance” is not a relevant consideration in our chosen semantic similarity measure whereas distance is relevant in the machine learning methods we consider. We find that the notion of distance introduces a bias in prediction results, similar to biases found in some semantic similarity measures (Kulmanov and Hoehndorf, 2017*a*; Cornish et al., 2018); using these methods should consequently be considered carefully, in particular as their blackbox nature makes it challenging to identify the reason for a prediction.

We identified and tested the impact of different biases within phenotype-based methods for finding candidate genes. We found a general study bias where disease-associated genes (or genes whose human ortholog is disease-associated) have generally more, and more specific, annotations than non-associated genes, and this affects not only semantic similarity measures but also machine learning methods; even more concerning, supervised machine learning methods can exploit biases in the data to make accurate predictions based on non-biological properties of the data (such as number and type of phenotype annotations). Again, use of blackbox models such as neural networks presents the danger of hiding the biases and how they are utilized in decision making.

The use of model organisms for understanding genotype– phenotype relations in humans is well established as a valuable strategy. Few organisms present a complete model of the human (Sundberg et al., 2013), but aspects of, for example, a disease phenotype might be studied more conveniently and with good fidelity in certain species or strains, yielding valuable insights from several MOs (Hmeljak and Justice, 2019). While the mouse is anatomically and physiologically closest to humans and therefore phenotypes are more likely to be easily related, we know that in particular systems or metabolic pathways valuable information can come from much more distant organisms, for example insights into cell cycle control or ageing from yeast (Pardo and Boriek, 2020). The computational use of model organism phenotypes to identify the underlying genetics of disease is a recent development, based on the premise that, as we do not have functional information for all genes in humans, combining this knowledge from model organisms can massively increase the knowledge that can be brought to bear (Mungall et al., 2010). Based on our data, there are 13,789 human genes that have an ortholog in the model organisms we investigate which has been assigned one or more phenotypes (11,672 human genes have an ortholog in the mouse with phenotype annotations; 3,418 genes in the fish; 6,462 in the fruitfly; and 1,871 in yeast), and therefore over 63% of human genes have orthologs in model organisms with phenotype annotations (Willyard, 2018). Figure 2 shows the pairwise overlap of genes with phenotypes in mouse, fish, fly and yeast.

**Fig. 2.**
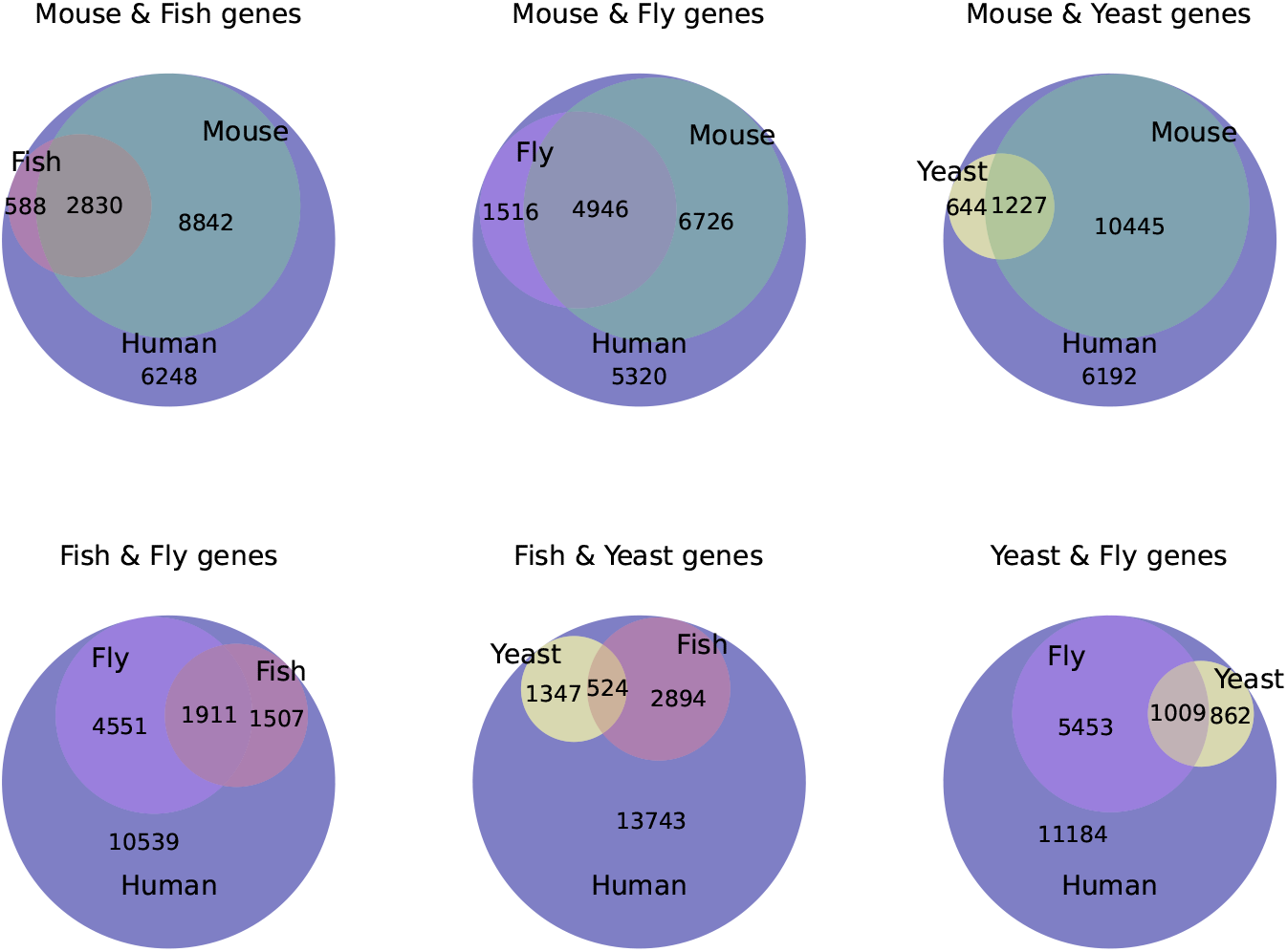
This figure present human genes with model organism orthologs. The pairwise intersection of model organisms with phenotypes is illustrated in sub-graphs. Each of these sub-graphs has 18508 human genes in total.

A question not so far addressed is which of these annotations add to the power of computational approaches to discover disease-associated genes. By using two cross-species phenotype ontologies we are able to show that the data from the mouse explains the majority of the human disease gene associations and that very little, if any, data from other models organisms contributes to this task. A previous focused study comparing mouse and fish phenotypes to predict disease genes in different disease categories for the Phenodigm algorithm (Oellrich et al., 2014) also showed the mouse to be overall more useful but suggested that the zebrafish made contributions in specific disease areas. The authors suggested that this may be due to increased coverage of cardiovascular diseases in the data from mutant fish. Our findings are consistent with this, and also suggest that, as a consequence, the resulting performance is due to biases in the number and specificity of annotations and not due to the intrinsic relatedness between phenotypes. An increased number of, and more specific, annotations will bias estimates of semantic similarity. The effect of these biases has been demonstrated when comparing between model organism and human disease phenotypes (Kulmanov and Hoehndorf, 2017*b*), and also when predicting gene–disease associations where this bias can be corrected when detected (Cornish et al., 2018).

We further demonstrate that assessment of contribution to disease gene identification depends critically on the methods used, and we present evidence that supervised machine learning methods systematically overestimate the contribution of model organisms, mainly by exploiting biases in phenotype data. Similar biases affect the evaluation of gene–disease and variant pathogenicity prediction methods (Grimm et al., 2015), and are challenging to detect and correct for. In the future, evaluation datasets and methods need to be developed that are less likely to overfit to biases in training and testing data, generalize well across different organisms, and are robust to noise in phenotype annotations.

## MATERIALS AND METHODS

### Ontologies

Several foundational ontologies are used for the axiomatisation of species-specific phenotype ontologies, and we reused them for the construction of the Pheno-e ontology. We used the Gene Ontology (GO) (Ashburner et al., 2000) downloaded from http://purl.obolibrary.org/obo/go.owl ; the Cell Ontology (CL) (Diehl et al., 2016) downloaded from http://purl.obolibrary.org/obo/cl-basic. owl; Phenotype and Trait Ontology (PATO) (Gkoutos et al., 2005) downloaded from http://purl.obolibrary.org/obo/pato.owl; Uber Anatomy Ontology (UBERON) (Mungall et al., 2012) downloaded from http://purl.obolibrary.org/obo/uberon.owl; Zebrafish Anatomy and Development Ontology (ZFA) (Van Slyke et al., 2014) downloaded from http://purl.obolibrary.org/obo/zfa.owl; Neuro Behavior Ontology (NBO) (Gkoutos et al., 2012) downloaded from http://purl.obolibrary.org/obo/nbo.owl; Biological Spatial Ontology (BSPO) (Dahdul et al., 2014) downloaded from http://purl.obolibrary.org/obo/bspo.owl; *Drosophila* Gross Anatomy Ontology (FB-BT) (Costa et al., 2013) downloaded from http://purl.obolibrary.org/obo/fbbt.owl.

The phenotype ontologies used were: Mammalian Phenotype Ontology (MP) (Smith, Goldsmith and Eppig, 2005) downloaded from http://purl.obolibrary.org/obo/mp.owl; Human Phenotype Ontology (HP) (Köhler et al., 2018) downloaded from http://purl.obolibrary.org/obo/hp.owl; Drosophila Phenotype Ontology (DPO) (Osumi-Sutherland et al., 2013) downloaded form http://purl.obolibrary.org/obo/dpo.owl and Fission Yeast Phenotype Ontology (FYPO) (Harris et al., 2013) downloaded from http://purl.obolibrary.org/obo/fypo.owl. The latest version of the ontologies is used for every update of Pheno-e; the results reported here use ontologies downloaded in February 2021.

### Data Sets and Phenotype Annotations

For constructing the model organism phenotype classes we used the following:

- From the ontologies MP (Smith, Goldsmith and Eppig, 2005), HP (Köhler et al., 2018), DPO (Osumi-Sutherland et al., 2013) and FYPO (Harris et al., 2013), we reconstructed the phenotype classes for mice, human, fly and yeast respectively.
- From FlyBase (Thurmond et al., 2018) we used `allele_phenotypic_data_fb_2021_01.tsv`, which provides the alleles phenotypes association using controlled vocabulary for *Drosophila melanogaster*. We used this file to create the abnormal anatomy classes (FBabAB).
- From ZFIN, we used `phenoGeneCleanData_fish.txt` which contains zebrafish gene–phenotype associations to create classes representing zebrafish phenotypes.

For generating representations of genes and diseases and for predicting gene–disease associations we used the following files downloaded on 07-Feb-2021:

- Human disease–phenotype annotations were obtained from the HPO database (Köhler et al., 2018) `phenotype_annotation.tab`. comprising manual and semi-automated annotations representing disease identifiers from three databases OMIM (Amberger and Hamosh, 2017), Orphanet (Weinreich et al., 2008) and DECIPHER (Firth et al., 2009).
- Mouse gene–phenotype annotations were obtained from the Mouse Genome Informatics (MGI) database (Ringwald et al., 2021) MGI_GenePheno.rpt which use MP (Smith and Eppig, 2012).
- We generated the fly gene–phenotype annotations using files from FlyBase (Thurmond et al., 2018), `allele_- phenotypic_data_fb_2021_01.tsv` which represents the allele–phenotype annotations and `fbal_to_fbgn_fb_- 2021_01.tsv` which contains allele–gene mappings.
- We obtained yeast gene–phenotype annotations from Pombase (Harris et al., 2013) which represent the phenotypes annotations for fission yeast *Schizosaccharomyces pombe* `phenotype_- annotations.pombase.phaf.gz`.

For evaluation purpose, we used two gene-disease association datasets from MGI (Ringwald et al., 2021). The first dataset is a “human” dataset that includes associations of human genes with Mendelian diseases using identifiers from OMIM database (*Online Mendelian Inheritance in Man (OMIM)*, 2020). This dataset includes 2,848 human gene, 3644 OMIM disease and 11,778 human gene–disease associations. The second dataset is a “mouse” evaluation set that includes associations of mouse genes with human disease and represents mouse models of human disease in the MGI database. This dataset contains 2,459 mouse genes, 2,157 disease and 8,101 mouse gene–disease associations. Both datasets are included in the file `MGI_DO.rpt` from MGI. The version we used was downloaded on Feb 2021.

To find orthologous genes between different organisms we used several files. Human–mouse orthology was obtained from `HMD_HumanPhenotype.rpt` from MGI. Human–zebrafish and mouse–zebrafish orthologs were obtained from `human_- orthos.txt` and `mouse_orthos.txt` from ZFIN. We obtained human–fly orthology from `dmel_human_orthologs_- disease_fb_2020_06.tsv` from FlyBase. Human–yeast orthologs were obtained from `pombeorthologs`. We obtained mouse–fly and mouse–yeast orthologs from OMA (Train et al., 2017).

### Pheno-e and integration of model organism phenotypes

The PhenomeNET Ontology was developed by utilizing existing phenotype ontology class descriptions and reformulating them according to a set of ontology design patterns so that different phenotype ontologies can be integrated (Hoehndorf et al., 2011). The uPheno ontology similarly establishes bridging axioms to connect phenotypes from different species-specific ontologies (Matentzoglu, Osumi-Sutherland, Balhoff, Bello, Bradford, Cardmody, Grove, Harris, Harris, Köhler et al., 2019). The current version of the PhenomeNET ontology does not contain classes for yeast and fly phenotypes while these two species are covered in uPheno. We therefore expanded PhenomeNET to include phenotypes from fly and yeast. We obtained the phenotype class descriptions from the DPO and FYPO ontologies, and reformulated them using the PhenomeNET design patterns. The new classes we created use the pattern

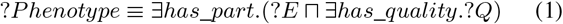

In this pattern, ?*E* characterizes the entity underlying the phenotype (either from an anatomy ontology or the GO) and ?*Q* is a quality from the PATO ontology. We use relations from the OBO Relation Ontology (Smith, Ceusters, Klagges, Köhler, Kumar, Lomax, Mungall, Neuhaus, Rector and Rosse, 2005); the relations we use in constructing PhenomeNET and Pheno-e include *part-of, results-from, during, has-quality, has-central-participant, occurs-in*, and *towards*.

FlyBase has two types of abnormal phenotype classes; it associates alleles with classes from the DPO as well as with classes from the fly anatomy ontology, indicating that an anatomical or developmental structure was found to be abnormal in a mutant fly. In order to integrate those anatomical abnormalities in the PhenomeNET ontology and therefore use them in cross-species phenotype analysis, we added the abnormal anatomical structures as new classes in PhenomeNET and associated the alleles with these classes:

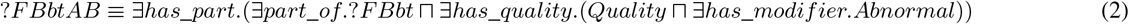

*Quality* and *Abnormal* are classes from the PATO ontology. For example, `FBal0148512` is an allele associated with wing abnormalities (Végh and Basler, 2003), and we associate the allele with the newly defined class *Phenotypic abnormality of wing*, defined accordingly to the pattern in 2 where ?*FBbt* is the class *Wing* from the fly anatomy ontology.

Similarly to PhenomeNET, we define homologous and analogous anatomical structures as equivalent (for the purpose of the ontology). For example, we defined the nervous system in fly (`FBbt:00005093`) to be equivalent to the nervous system in the zebrafish (`ZFA:0000396`), and the nervous system in the Uberon multi-species anatomy ontology (`UBERON:0001016`). Through these equivalence class assertions, we can deductively infer an equivalence between *nervous system phenotype* (`MP:0003631`), *abnormal neuroanatomy* (`FBcv:0000435`), and *Abnormality of the nervous system* (`HP:0000707`), thereby enabling the direct comparison of mouse, fly, and human phenotypes.

The extended PhenomeNET ontology (Pheno-e) contains 16,083 human phenotype (HP) classes, 13,698 mammalian phenotype (MP) classes, 35,954 Zebrafish phenotype classes (PHENO classes, defined in Pheno-e), 3,111 fly phenotype classes (FBcv classes and abnormal anatomy FBbtAB classes), and 7,636 classes of yeast phenotype (FYPO) classes.

We use phenotype datasets consisting of 8,031 OMIM diseases annotated with HP classes, 14,210 mouse genes annotated with MP classes, 6,182 zebrafish genes annotated with PHENO classes, 13,512 fly genes annotated with FBbtAB and FBcv classes, and 4,443 yeast genes annotated with FYPO classes.

Using automated reasoning over the Pheno-e ontology, we are able to infer relations between classes from different organisms; in particular, we are able to automatically infer whether two classes are equivalent or whether one class is a subclass of another class. We show the number of inferred relations in Pheno-e between the different species in Table 5, and for uPheno in Table 6. The tables show that it is possible to relate a large number of model organism phenotypes to human phenotypes through the Pheno-e and uPheno ontologies.

**Table 5.** Pheno-e summary of direct and indirect inferred sub-classes and super-classes axioms between different organisms phenotype classes

| Subclass Matrix |  |  |  |  |  |
| --- | --- | --- | --- | --- | --- |
|  | Human | Mammalian | Zebrafish | Drosophila | Yeast |
| Human Phenotypes | 5484 | 2086 | 1189 | 79 | 54 |
| Mammalian phenotypes | 2669 | 5806 | 2132 | 98 | 110 |
| Zebrafish phenotypes | 4016 | 6155 | 17617 | 398 | 937 |
| Drosophila phenotypes | 120 | 173 | 209 | 4143 | 28 |
| Fission yeast phenotpes | 62 | 128 | 332 | 25 | 2457 |

| Superclass Matrix |  |  |  |  |  |
| --- | --- | --- | --- | --- | --- |
|  | Human | Mammalian | Zebrafish | Drosophila | Yeast |
| Human Phenotypes | 15795 | 15389 | 15573 | 15573 | 7804 |
| Mammalian phenotypes | 13166 | 14096 | 13160 | 13160 | 7797 |
| Zebrafish phenotypes | 35954 | 30793 | 35951 | 35951 | 17916 |
| Drosophila phenotypes | 3017 | 2530 | 3016 | 19153 | 153 |
| Fission yeast phenotpes | 5807 | 3987 | 5807 | 5807 | 7322 |

**Table 6.** uPheno summary of direct and indirect inferred shared ancestor generic class count between different organisms phenotype classes

| Shared Ancestor Generic class count |  |  |  |  |  |
| --- | --- | --- | --- | --- | --- |
|  | Human | Mammalian | Zebrafish | Drosophila | Yeast |
| Human Phenotypes | 6414 | 2903 | 1426 | 91 | 67 |
| Mammalian phenotypes | 2903 | 7786 | 2007 | 87 | 75 |
| Zebrafish phenotypes | 1426 | 2007 | 6446 | 139 | 142 |
| Drosophila phenotypes | 91 | 87 | 139 | 177 | 53 |
| Fission yeast phenotpes | 67 | 75 | 142 | 53 | 177 |

Figure 3 illustrates an example of inferred relations between phenotype classes of different organisms and resources. In this example, the class *Abnormal T cell activation* (`HP:0410035`) has as a (zebrafish) superclass *Phenotypic abnormality of cellular process* (`PHENO:32859`). This inference was made because of the background available from PATO and GO as the class *process quality* (`PATO:0001236`) is a subclass of *quality* (`PATO:0000001`), and *T cell activation* (`GO:0042110`) is a subclass of *cellular process* (`GO:0009987`).

**Fig. 3.**
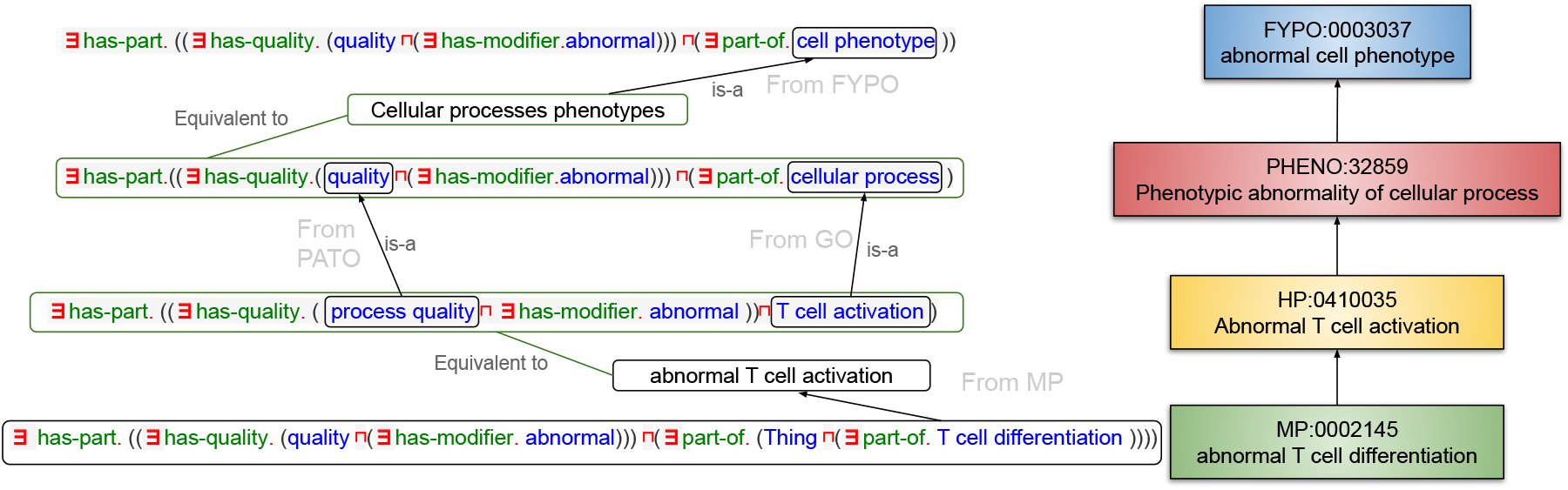
Example of inferred hierarchy relating classes from different organisms phenotypes.

### Phenotype similarity

We apply a set of different methods to compare the similarity of phenotypes associated with a loss of function model organism mutant and human disease phenotypes.

#### Resnik semantic similarity

We calculated Resnik similarity (Resnik, 1995) between genes and diseases annotated with phenotype classes; the use of integrated phenotype ontologies enables the direct comparison of phenotypes.

Resnik’s similarity is a similarity measure based on information content, defined as

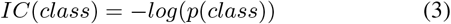

where the probability of a class is defined as the frequency of annotation with the class. The similarity between two ontology classes is defined as the information content of the most informative common ancestor (MICA) of two classes:

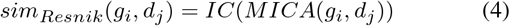

As we compare groups of classes, we use the best match average method (BMA) (Pesquita et al., 2009) to calculate the similarity between genes and diseases:

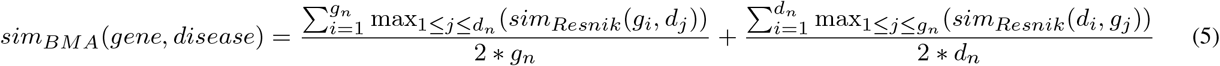

#### OPA2Vec

As second method, we “embed” phenotypes in a real-valued vector space using OPA2Vec (Smaili et al., 2018*a*); an embedding is a structure-preserving map from one algebraic structure (ontology axioms) into another (vector space), i.e., an embedding preserves (some) properties of the first structure within the second. OPA2Vec is mainly based on preserving syntactic relations in asserted and inferred ontology axioms.

We generated embeddings for diseases and genes using OPA2Vec based on the phenotypes associated with the diseases and genes and the axioms in the phenotype ontology. For the training, we use the skip gram, set `mincount` to 0, embedding size to 100, and window size to 5. We then compute similarity between genes and diseases based on the cosine similarity of their embeddings.

#### DL2Vec and OWL2Vec*

Another method we used to generate feature embeddings is the DL2Vec (Chen et al., 2020) method. DL2Vec converts description logic axioms into an undirected graph representations and uses a random walk to explore the graph; the walks are then treated as sentences and encoded using a language model. The graph is generated from the ontology axioms, and each phenotype class becomes one node in this graph; we add the gene and disease identifiers to this graph and connect them to the phenotype classes with which they are annotated. To generate the walks we choose the walk length to be 30, with 50 number of walks, and we used a skip gram method with window size set to 10, `mincount` to 1, and embedding size 100.

OWL2Vec*(Chen et al., 2021) is an embedding method similar to DL2Vec and based on a similar graph representation. OWL2Vec* graphs are directed and do not include equivalence or disjoint class axioms. We use random walker with walk depth 7 and 30 iterations with projection on structure document. and we used a skip gram method with window size set to 5, `mincount` to 1, and negatives to 5 and embedding size 100.

For both DL2Vec and OWL2Vec* embeddings, we compare the phenotypic similarity between genes and diseases using cosine similarity.

#### Prediction of gene–disease associations

In addition to predicting gene–disease associations based on phenotypic similarity, we also use supervised prediction of these associations. For this purpose, we use a multilayer perceptron (MLP) with a single hidden layer. The input of the MLP is the concatenated embeddings of a disease and a gene. We use a hidden layer half the size of the input and a binary output using a sigmoid function, indicating whether the gene and disease are associated through a gene–disease relation or not; we further use the value of the sigmoid to rank genes for a disease. We randomly generated five negatives to each positive. For the training, we use the Adam optimizer (Kingma and Ba, 2014) with a learning rate of 0.001 and maximum number of iterations of 300. To evaluate, we used 10-fold cross validation, stratified by diseases.

#### Naïve classifier

We hypothesize that some of our results are due to imbalanced or biased data. To test this hypothesis, we define a “naïve” classifier that predicts gene-disease associations on the basis of the information content of phenotypes of a gene alone; this “classifier” ranks all genes based on the sum of the information content of their phenotype annotations, sorts genes in descending order, and predicts the same ranked list of genes for each disease (i.e., the classifier is independent of the disease). The aim of this “naïve” classifier is to test whether genes annotated with more and more specific phenotypes are generally more likely to be associated with a disease, i.e., it tests for a kind of annotation bias.

### Evaluating predictive performance

Our evaluation is based on estimating how well the different approaches rank disease-associated genes given a set of disease-associated phenotypes, for phenotypes from different organisms. Higher phenotypic similarity between a gene and a disease indicates higher likelihood that the gene (or its human ortholog) is associated with that disease. We evaluated two data sets from the MGI file `MGI_DO.rpt`, one for human gene–disease associations from OMIM and another MGI-curated dataset of mouse models of human disease.

For the evaluation, for each disease *D*_*i*_ in our evaluation set, we rank all genes *G*_1_, …, *G*_*n*_ based on their phenotypic similarity to *D*_*i*_. For each disease *D*_*i*_, we determine the rank (or ranks) at which the associated gene (or genes) appear in this ranked list. We use this information to determine the false positive and true positive rate at each rank; we average the true and false positive rates across all diseases and use this to determine the receiver operating characteristic (ROC) curve and the area under the ROC curve (ROCAUC).

When using supervised methods to predict gene–disease associations, we use the same evaluation in a 10-fold cross validation setting, and we rank genes based on the output of the sigmoid unit of our machine learning model.

### Implementation

We used several tools and libraries, such as the OWLAPI for generating the ontology groovy and python scripts for data processing. We also used several python libraries like sklearn, numpy, pandas, PyTorch (Paszke et al., 2017) for the supervised learning. For calculating Resnik semantic similarity we used the Semantic Measures Library (SML) (Harispe et al., 2013).

## Acknowledgements

We acknowledge use of resources from the KAUST Supercomputing Laboratory. PNS acknowledges the support of The Alan Turing Institute.

## Contribution

PNS and RH conceived of the experiments; SMA, PNS and RH designed and interpreted the experiments; SMA performed and implemented all computational and statistical experiments; PNS and RH acquired the funding.

## Data availability

All data and software required to reproduce our results are freely available at https://github.com/bio-ontology-research-group/Module_organism_phenotypes

